# Protein-specific signal peptides for mammalian vector engineering

**DOI:** 10.1101/2023.03.14.532380

**Authors:** Pamela O’Neill, Rajesh K Mistry, Adam J. Brown, David C. James

## Abstract

Expression of recombinant proteins in mammalian cell factories relies on synthetic assemblies of genetic parts to optimally control flux through the product biosynthetic pathway. In comparison to other genetic part-types, there is a relative paucity of characterized signal peptide components, particularly for mammalian cell contexts. In this study, we describe a toolkit of signal peptide elements, created using bioinformatics-led and synthetic design approaches, that can be utilized to enhance production of biopharmaceutical proteins in Chinese Hamster Ovary cell factories. We demonstrate, for the first time in a mammalian cell context, that machine learning can be used to predict how discrete signal peptide elements will perform when utilized to drive ER translocation of specific single chain protein products. For more complex molecular formats, such as multichain monoclonal antibodies, we describe how a combination of *in silico* and targeted design rule-based *in vitro* testing can be employed to rapidly identify product-specific signal peptide solutions from minimal screening spaces. The utility of this technology is validated by deriving vector designs that increase product titers ≥ 1.8x, compared to standard industry systems, for a range of products, including a difficult-to-express monoclonal antibody. The availability of a vastly expanded toolbox of characterized signal peptide parts, combined with streamlined *in silico/in vitro* testing processes, will permit efficient expression vector re-design to maximize titers of both simple and complex protein products.

**GRAPHICAL ABSTRACT:** 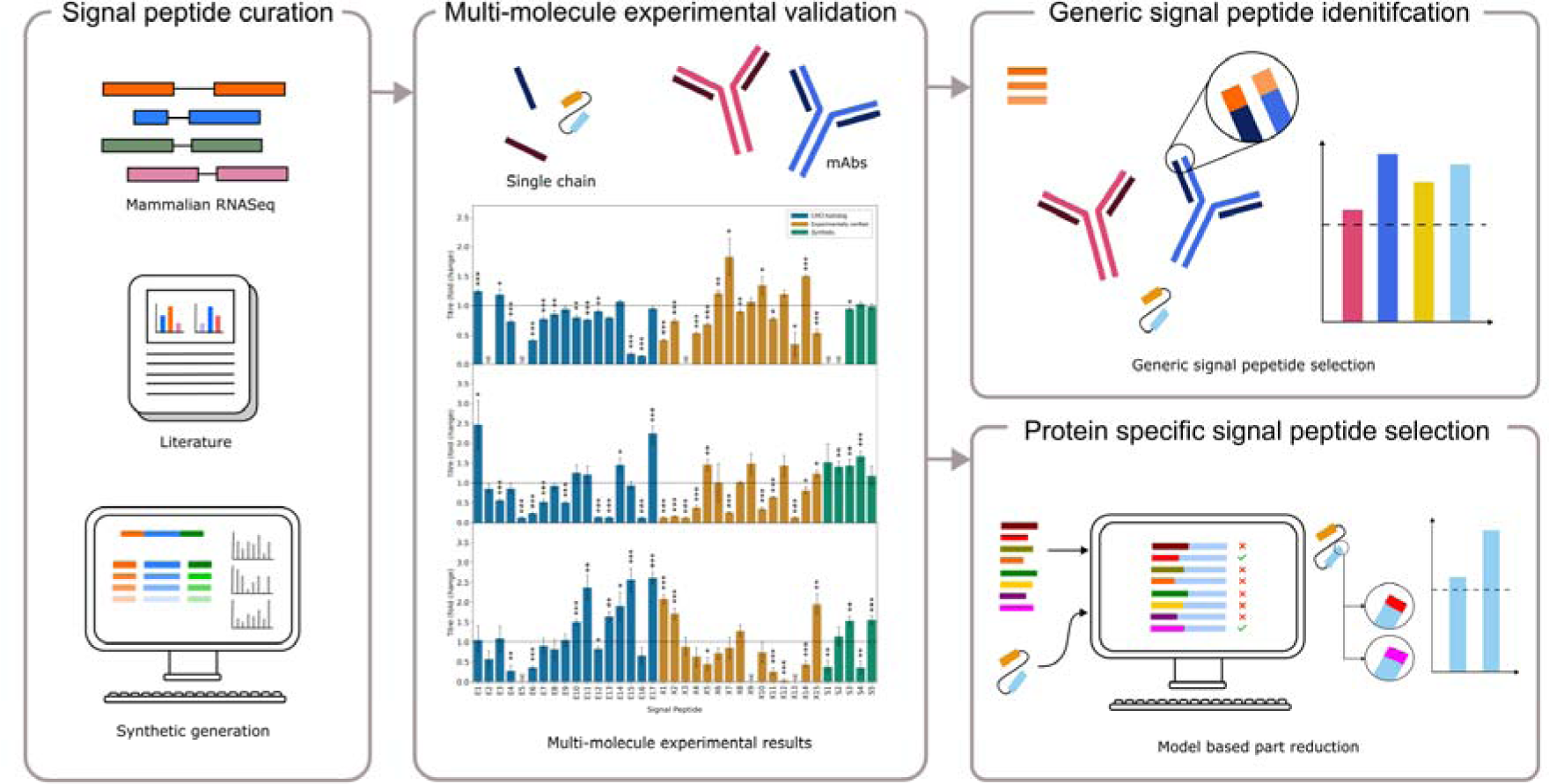

## 1. INTRODUCTION

Recombinant proteins are the principal molecular format of biopharmaceutical products, where Chinese hamster ovary (CHO) cells are a dominant cell factory utilized for their biomanufacturing. Monoclonal antibodies (mAbs) are the most common product-type in development, representing 53.5% of all biopharmaceutical approvals between 2018-2022 (1). Over the past two decades significant titer increases have been achieved via cell (2), vector (3), process (4) and media (5) engineering. Despite significant advances in biomanufacturing system outputs, new technologies are critically required to enhance production of increasingly complex product formats, such as fusion proteins and tri-specific mAbs (6). These proteins are commonly referred to as ‘difficult-to-express’ owing to low product yields, where process optimization is time and cost-intensive (7-9).

Advances in the synthetic biology field, particularly in DNA sequence engineering, have significantly expanded opportunities to improve CHO cell expression vector design (10). While some vector components have been widely studied, with associated availability of DNA part libraries (11), mammalian signal peptides remain relatively unexplored. As all recombinant protein products are secreted from the host cell factory, they have an absolute requirement to be paired with an appropriate signal peptide, a short N-terminal amino acid sequence that facilitates co-translational translocation of nascent polypeptides into the endoplasmic reticulum (12, 13). While signal peptides adhere to a generic three-domain structure, comprising a basic N-domain, a hydrophobic H-domain and a slightly polar C-domain (14), the sequence features underpinning their performance are relatively poorly understood. Recent studies have begun to elucidate mechanistic design rules that govern whether a signal peptide will generally encode low or high ER translocation rates (15), however (i) this work is predominantly in bacterial systems (16, 17), and (ii) there is a paucity of information regarding context-specific functionality, whereby individual signal peptide performance is highly variable dependent on the partner-protein used.

Previous work in CHO cells has identified bottlenecks in the secretory pathway as a limiting factor in production yields (18-20). Multiple studies have determined that the rate of ER translocation is a critical control parameter in the product biosynthetic pathway, where utilization of novel signal peptides has been shown to significantly enhance the titer of a wide range of recombinant proteins (21). Although screening small panels of signal peptide parts typically identifies a component that permits increased system output, as compared to common industry-used sequences, protein-partner specificity necessitates trial and error screening which is intractable in time-sensitive applications such as biopharmaceutical cell line development processes. The introduction of predictive tools based on machine learning (ML) and deep learning, such as SignalP (22), have provided a streamlined approach to identifying and selecting signal peptides which are likely to facilitate correct peptide cleavage. Moreover, ML approaches have recently been employed to create tools that can create (16) and select (17) signal peptides to function within specific protein-partner contexts. However, such tools are not available for mammalian cell contexts, nor have they been described for situations where multiple polypeptide chains need to be simultaneously expressed, as is the case for mAb LC and HC molecules.

In this study, we have employed three distinct design routes to create a library of signal peptide components for use in CHO cell vector engineering. We validated the utility of this toolbox, the largest panel of signal peptides ever designed and tested in CHO cell systems, by using it to identify parts that facilitated significant titer increases (compared to standard industrial components) for a range of protein products. Critically, we also describe *in silico* ML-based and design rule-based *in vitro* screening methods that substantially reduce the testing required to identify product-specific signal peptides for both simple single-chain molecules and complex multi-chain proteins. This technology can be applied to rapidly derive synthetic signal peptide-protein partner assemblies that optimize ER translocation rates to enhance outputs from biopharmaceutical manufacturing systems.

## 2. RESULTS AND DISCUSSION

### 2.1 Creating a synthetic signal peptide toolkit for mammalian host cell expression vector engineering

A library of 37 signal peptides was designed, containing 17 CHO homologous, 15 experimentally-verified, and 5 synthetically designed signal peptides (Table 1). The purpose of these sub-groups was to assess a broad range of signal peptides and their impact on transient protein expression in a CHO-K1 derived host. While synthetic constructs have the potential to move significantly beyond the performance of naturally evolved sequences (3, 11), the design rules underpinning signal peptide functionality are relatively poorly-understood, resulting in poor predictability of synthetic element activity. Accordingly, preference was given to experimentally verified sequences, and CHO homologous signal peptides, based on the hypothesis that endogenous parts may exhibit optimized interactions with the CHO cell factory translocation machinery.

**Table 1:**
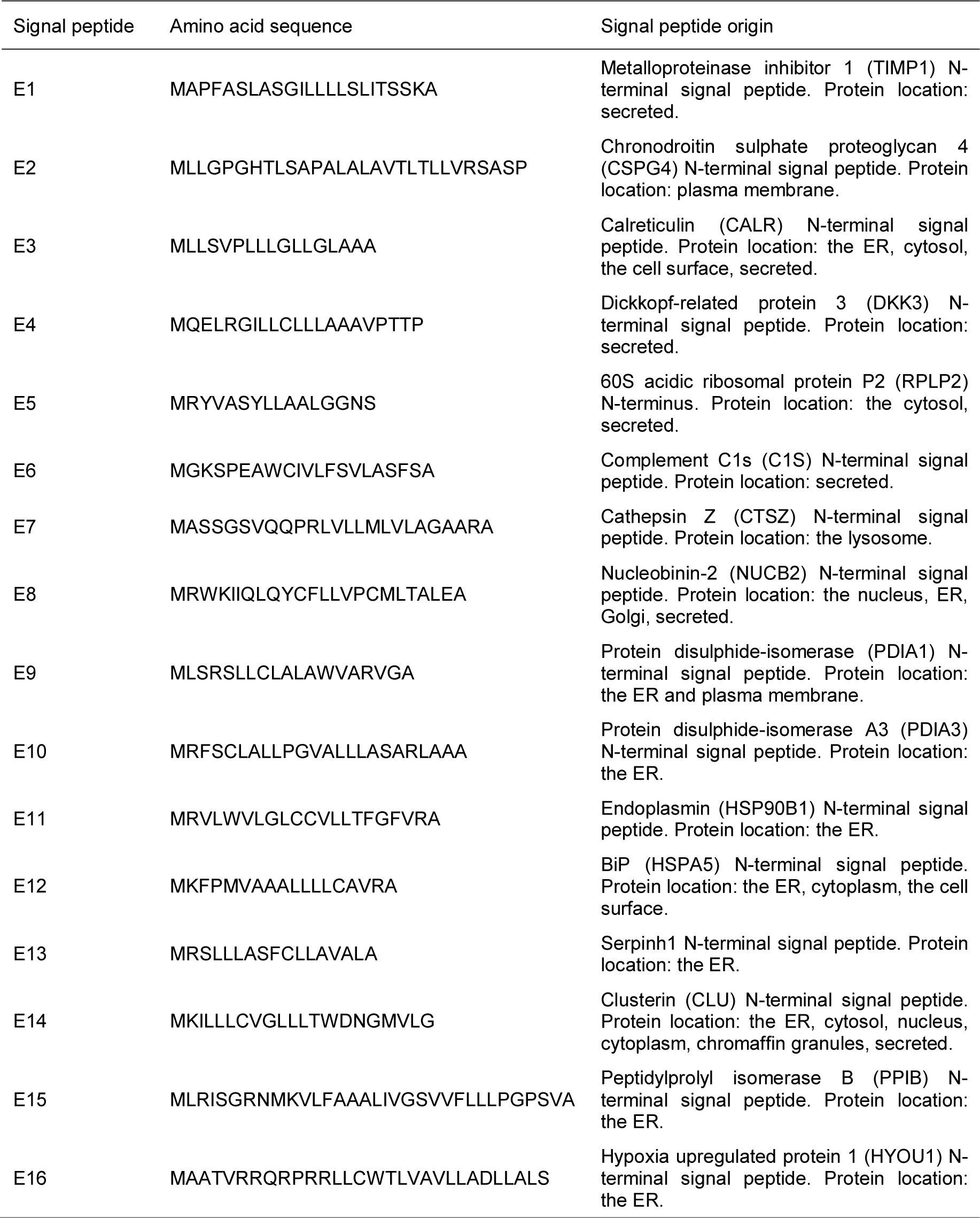
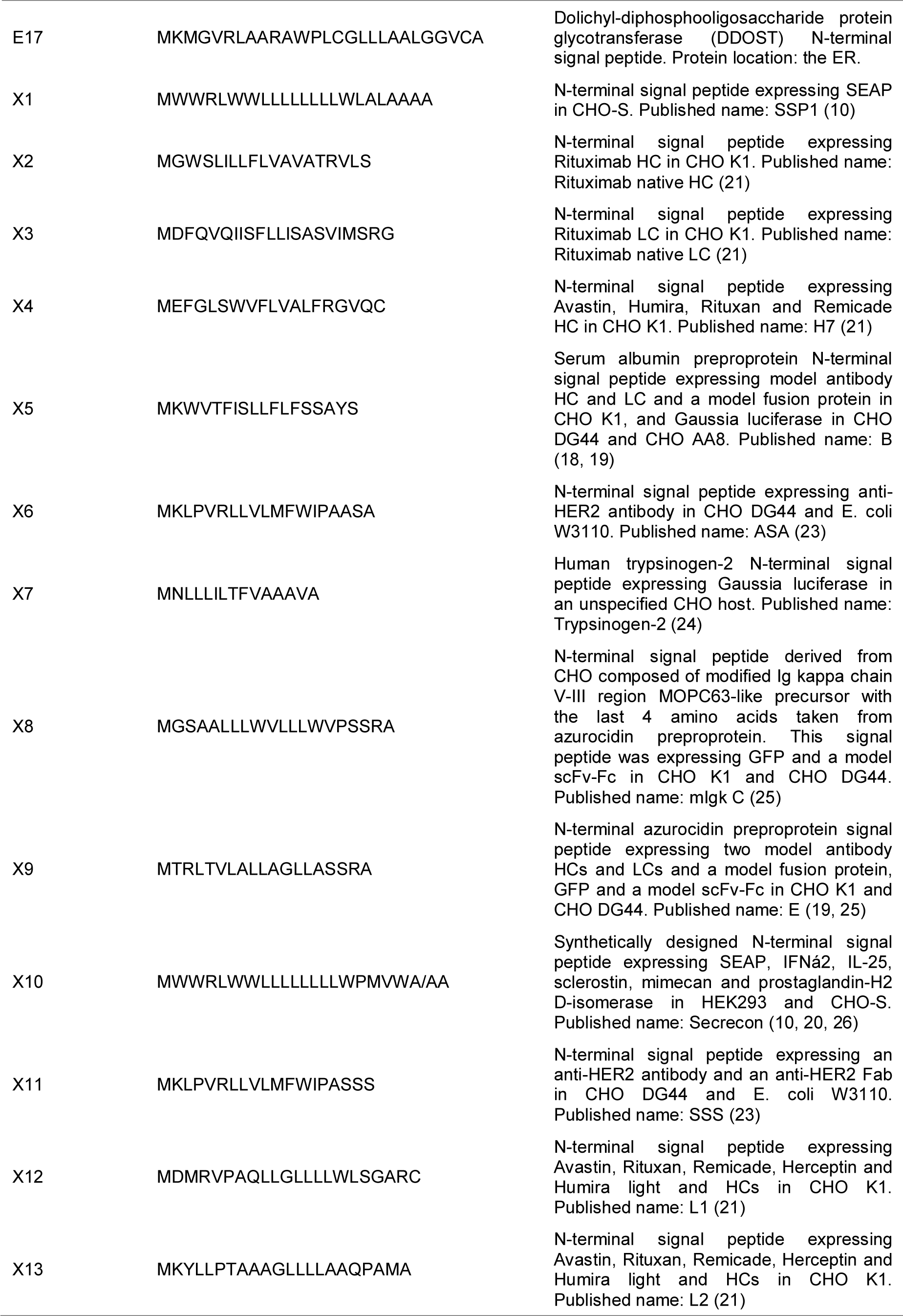
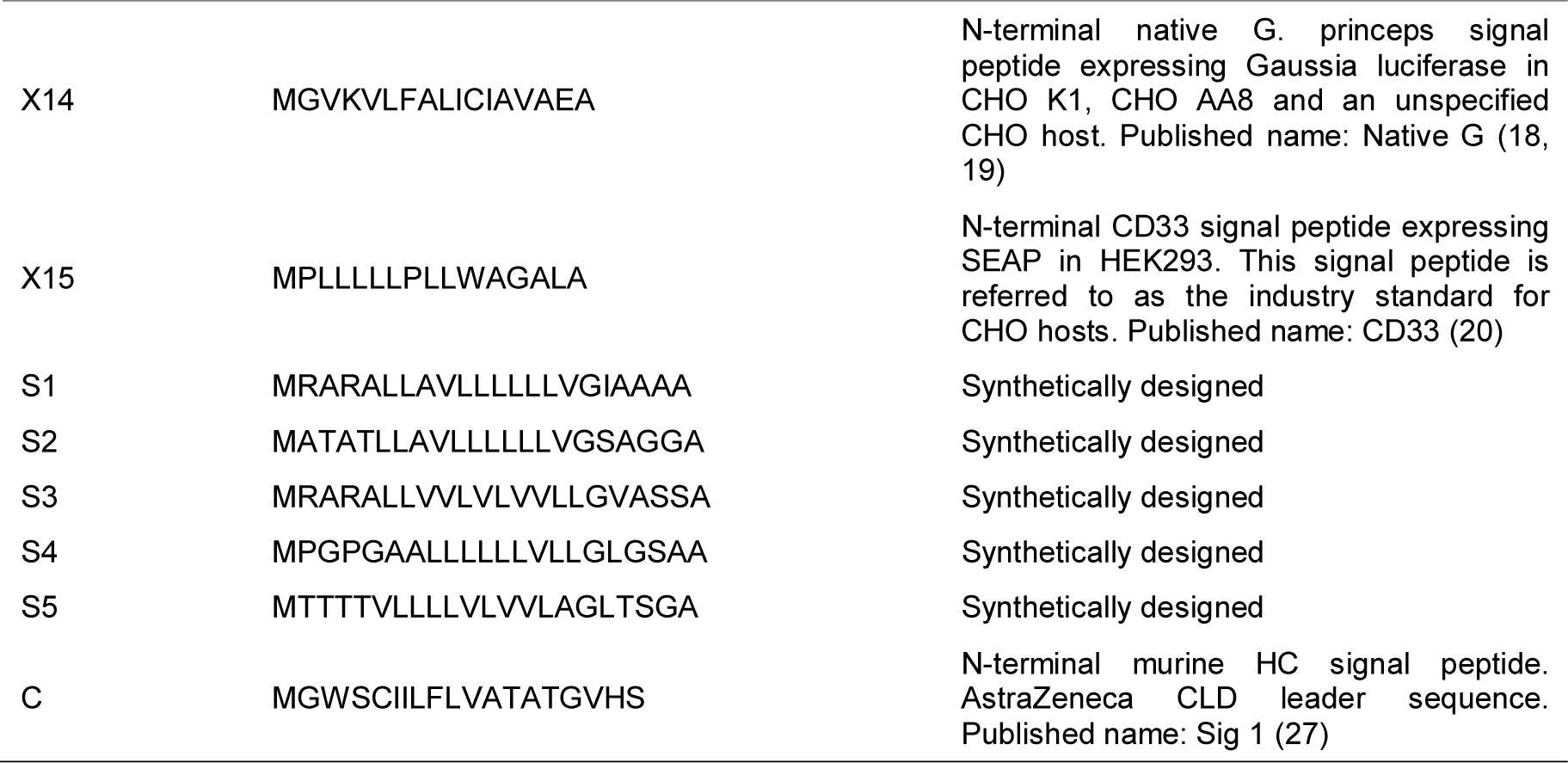
Origin and amino acid composition of 37 signal peptides used to construct ScfV, ETE and DTE mAb expression plasmids. Signal peptide ‘C’ is used as an industrially relevant standard reference signal peptide (ISC). E = CHO homologous, X = experimentally verified, S = synthetically designed.

CHO homologous signal peptides were selected from an in-house RNASeq dataset that profiled the transcriptome of a mAb producing CHO-S cell line. Proteomic datasets were not utilized due to the relatively low coverage typically obtained in CHO cell proteomic studies, which would have significantly restricted the design space. This dataset was first ranked by mRNA abundance (FPKM) then filtered using SignalP4.1 to only show proteins predicted to contain an N-terminal signal peptide. The 17 signal peptide-containing proteins with highest mRNA abundance were selected for *in vitro* testing, based on the hypothesis that very highly expressed genes may encode signal peptides that facilitate a relatively high translocation rate. It is of note that 11 of the 17 signal peptides taken from this dataset are present in the endoplasmic reticulum (ER), with three of these 11 proteins also being secreted from the cell. Of the remaining six signal peptides, four (E1, E4, E5, E6) are taken from secreted native proteins and two (E2, E7) from native proteins that are only located in the plasma membrane and the lysosome respectively. Signal peptides of particular interest from the CHO homologous group were those taken from ER chaperone proteins (E11: endoplasmin, E12: BiP), proteins involved in ER protein processing (E9: PDIA1, E10: PDIA3, E15: PPIB, E17: DDOST) and proteins which are stress-induced (E16: HYOU1) as these pathways are upregulated in recombinant protein production (28). It was therefore hypothesized that the signal peptides from these proteins may be preferentially recognized and imported into the ER in protein-producing CHO cells.

Literature-mined signal peptides were selected based on the criteria that the signal peptide facilitated increased expression of at least one recombinant protein, in comparison to a control, in a CHO cell host. Following a comprehensive search of published studies testing signal peptide functionality in CHO cells, a total of 15 discrete signal peptide sequences were identified that met this selection criteria. Whilst all of these constructs have the ability to drive high levels of ER translocation in a CHO cell context, due to protein partner specificity we did not hypothesize that all of these sequences would perform well when combined with our recombinant product testing panel.

Synthetic signal peptides were created according to design rules that we previously applied when generating a genetic component assembly toolkit for CHO cells. Specifically, as higher-level signal peptide sequence features are poorly understood, we used a simple domain-based design to create novel sequences with defined N-, H-, and C-regions. A database of experimentally verified mammalian signal peptides were extracted from signalpeptide.de before separating each sequence into constituent domains. The average size of each domain was calculated (N = 5AA, H = 12AA, C = 5AA) and synthetic domain sequences were then created *in silico* as detailed in Section 4.1. Briefly, experimentally-verified signal peptide domains were analyzed to identify conserved amino acids in each region, which were then randomly assembled to create thousands of unique configurations. Additional design constraints were placed on synthetic C-domain creation, applying rules from literature that have been shown to enhance/facilitate the functionality of this region (14, 29, 30).

Combining synthetic domain sequences in all possible permutations resulted in a library of > 1×10^12^ signal peptide constructs. Levinshtein distancing analysis was performed to identify sequences with highest heterogeneity (ie. to find discrete points within the design space), resulting in 1.18×10^6^ signal peptides. These sequences were analyzed using SignalP, where 0.03% were predicted to be functional signal peptides (D score > 0.7). We note that as SignalP is trained using endogenous sequences, and synthetic constructs which move significantly beyond the natural design space therefore have an increased chance of being designated as non-functional signal peptides. To maximize the chances of identifying high-performing synthetic constructs, a panel of five sequences with D scores > 0.7 were selected for *in vitro* testing.

SignalP demonstrates a eukaryotic cleavage precision score of between 0.795-0.914 for cleavage predictions ±0AA -±3AA. Accordingly, previous experimental analyses (covering a range of signal peptides and polypeptides) clearly demonstrate the accuracy of cleavage site predication by SignalP (31). Moreover, previous analyses of mAbs with different signal peptides expressed by CHO cells, both in this laboratory (data not shown) and published by others (21), have shown that >98% of mAb polypeptides are correctly processed – strongly aligning with a SignalP cleavage prediction. As this study is focused on the effect of signal peptide sequence on recombinant protein production rate (and considering that the study reports the expression of 333 different signal peptide/protein combinations), we concluded that measurement of cleavage efficiency in all cases was impractical and that to measure only a small number would be inconclusive (ie. it cannot be concluded that a demonstration of correct cleavage for one signal peptide-protein combination infers that a different combination will be correctly processed). It is clear however that ultimately, use of a specific signal peptide-protein combination in a particular context such as biopharmaceutical production would require analytical confirmation of correct cleavage.

### 2.2 A rationally designed panel of signal peptides permits molecule-specific optimization of recombinant protein production in mammalian cells

To evaluate the performance of our designed signal peptide panel, each signal peptide was tested in combination with three industrially relevant biopharmaceutical proteins, an ScFv fusion protein, an easy to express (ETE) mAb and a difficult to express (DTE) mAb. The optimal ratio of mAb HC:LC protein expression is highly product-specific, and utilizing the same signal peptide for both chains is unlikely to permit maximal product titers. Accordingly, we first tested the signal peptide library in combination with the LC constructs alone (i.e., without co-expression of cognate HCs; LCs were chosen as they are secreted, permitting simple quantification). Vectors containing each signal peptide in combination with partner product coding sequences were transiently transfected into a CHO-K1 derived host cell line. Relative product titers were determined by ELISA (mAb LCs) or ValitaTitre™ (ScFv fusion) at the end of 5-day fed batch production processes. As shown in Figure 1, signal peptides within the toolbox exhibited variable performance, where for each product molecule the test elements enabled titers ranging from no expression (NE) to a ≥ 1.8x increase, relative to an industrial standard control construct (ISC). The maximum titer increase facilitated was variable between each product, where the best performing signal peptides enhanced yields by 1.8-fold (X7), 2.5-fold (E1) and 2.7-fold (E17) for the ETE LC, DTE LC and ScFv fusion product respectively. In each case, at least 6 signal peptide elements were identified that out-performed ISC. However, similar to previous studies (20, 21) the relative performance of each signal peptide was typically highly product-specific. Indeed, no signal peptide element facilitated titer increases across all three molecules, validating the use of a toolbox approach. Moreover, a pair-wise analysis of library function across all three test molecules showed that there were no significant correlations in signal peptide performance (Fig. 2A-C). There is limited mechanistic understanding of the rules governing how a signal peptide performs when in combination with a specific partner protein sequence. Although LC sequences share significant similarities in amino acid composition and physiochemical properties, our data align with previous studies showing that the functionality of a discrete signal peptide is variable across different mAb molecules (21). One notable difference in protein sequence between the LC molecules and the ScFv fusion protein is the presence of basic amino acid residues between position +1 -+10 in the latter. Basic amino acid residues have been shown to directly affect the function of different signal peptides dependent on their relative hydrophobicity and polarity (32, 33). However, while general hypotheses can be made as to why a panel of signal peptides exhibits variable performance between two different molecules, the relative sequence features underpinning this are poorly understood.

**Figure 1:**
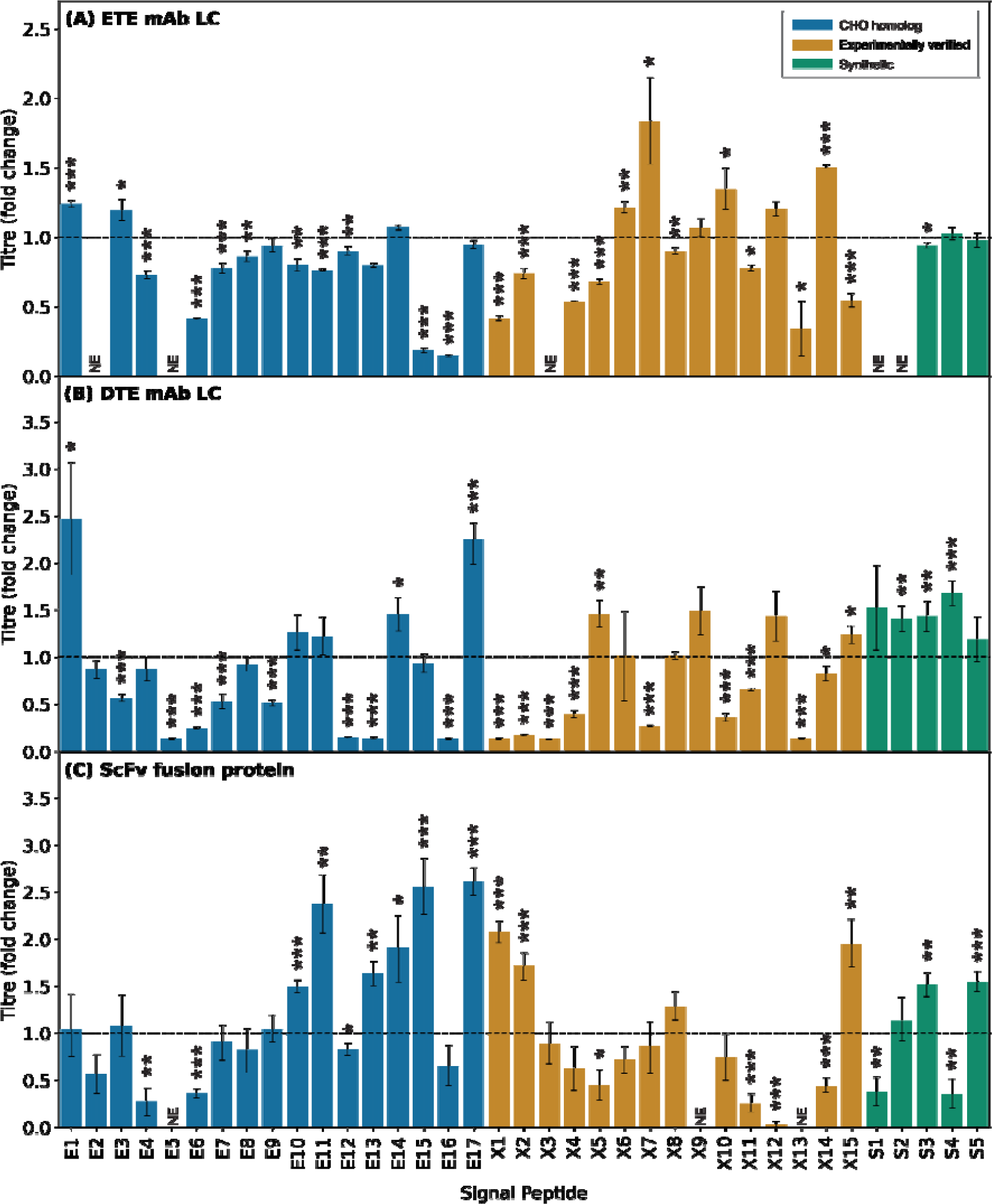
Choice of signal peptide significantly impacts production of single chain recombinant proteins. Expression constructs (a total of 111 unique constructs) each encoding one of 37 mammalian signal peptides (Table 1) with one of three recombinant single chain molecules (A, B, C) were independently transfected into CHO-K1 cells followed by measurement of secreted recombinant protein titer after 5d culture. (A) ETE (easy to express) IgG1 mAb LC, (B) DTE (difficult to express) IgG1 mAb LC and (C) ScFv fusion protein. Data were normalized with respect to the mean volumetric titer observed on transfection of the respective recombinant protein construct harboring a control murine Ig HC signal peptide -ISC (MGWSCIILFLVATATGVHS; (27), dotted line). Signal peptides are divided into three groups, E (CHO homologous, blue), X (literature-mined, gold) and S (synthetic, green); Table 1. NE denotes no measured expression. Each bar shows the mean ± standard deviation derived from three independent transfections, each performed in duplicate. Statistical significance is defined as p ≤ 0.05 (* = p≤ 0.05, ** = p≤ 0.01, *** = p≤ 0.001).

**Figure 2:**
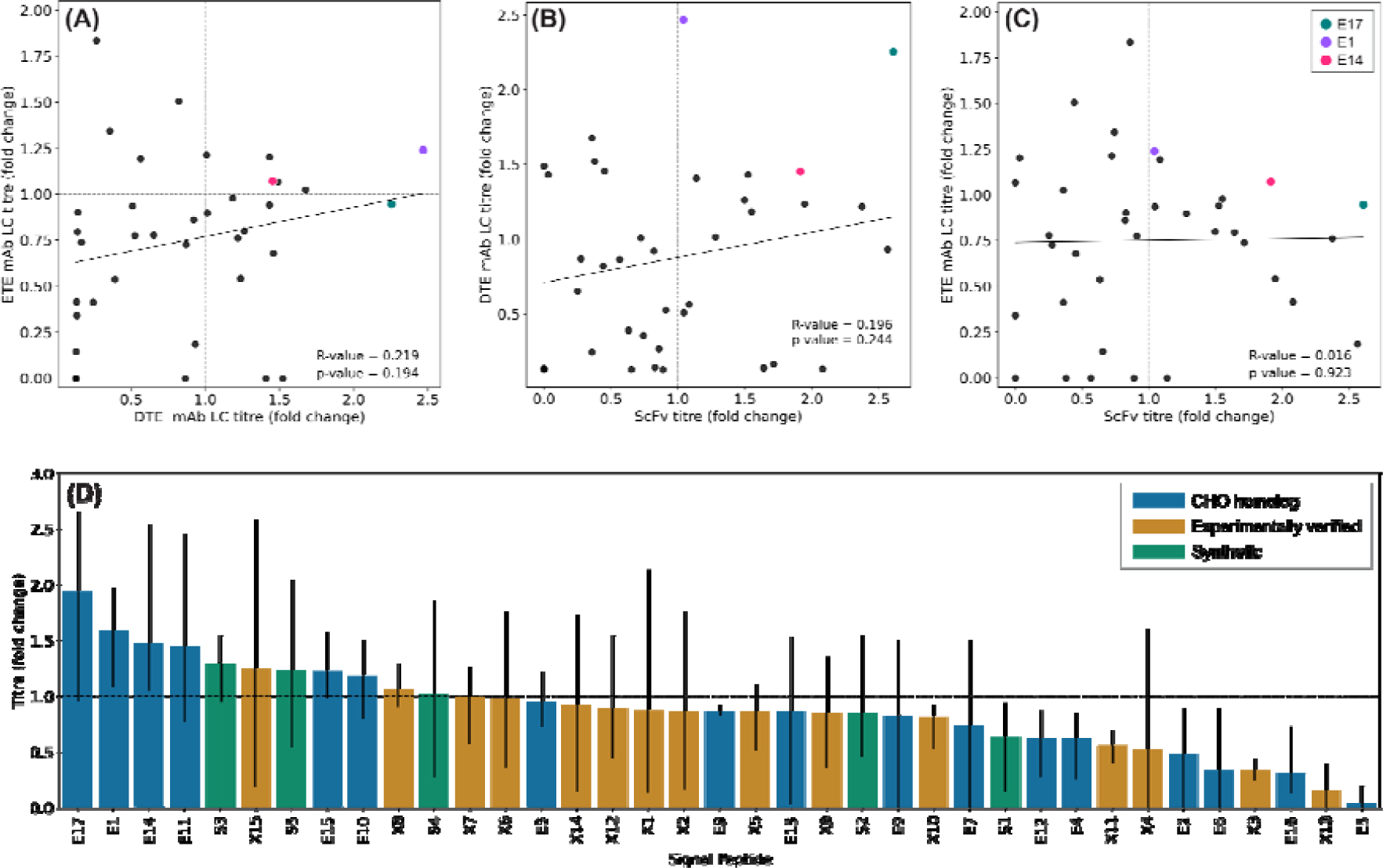
Effect of signal peptide on recombinant protein production is molecule specific. Derived from the data shown in Fig. 1, production of ETE mAb LC, DTE mAb LC and ScFv fusion protein mediated by different signal peptides are generally not correlated (A-C). Signal peptides highlighted (E17, E1, E14) show generic good performance. Grey dashed line represents quadrant separation. However, CHO endogenous signal peptides (E17, E14, E1, E11) yielded maximum volumetric titers. Bars represent the mean recombinant protein titer across the three recombinant proteins tested for each signal peptide (Fig. 1). Error bars represent the volumetric titer range across the three recombinant proteins tested for each signal peptide. Data are normalized with respect to the respective recombinant protein ISC (dotted line).

Although it is intractable to identify a universal signal peptide sequence that performs optimally across a product portfolio, we hypothesized that it may be possible to significantly reduce the testing space required to select high-performing elements. The simplest classification within our signal peptide library is the method underpinning their design/selection, i.e., experimentally verified, CHO homologous (identified via bioinformatics analysis) and synthetically designed. As shown in Figure 1, none of these design routes were generally superior, where each group contained a mixture of low-high performing elements across each molecule. Moreover, the high-performing constructs for each product (i.e., those permitting increased yields compared to ISC) were not associated with a particular signal peptide type. We concluded that this validated our initial toolbox strategy to derive elements from various design pathways. Moreover, it indicates that the signal peptide design (i.e., synthetically designed elements) and selection (i.e., bioinformatics-derived) methods that we employed in this study should also be effective in other contexts (e.g., different cell-types).

It is perhaps surprising that constructs which have previously been experimentally verified as driving high levels of ER translocation did not generally exhibit more predictable function than our newly-identified elements (i.e. CHO homologous and synthetically designed signal peptides). This highlights a key advantage and disadvantage associated with signal peptide design/selection, namely that (i) the design space permits relatively simple identification of constructs that have enhanced performance compared to incumbent standards, but (ii) their functionality is typically highly context-specific, dependent on the associated product molecule. While the identification of multiple novel signal peptides that can be deployed to enhance product titers is a valuable resource, ideally the testing space would be minimized. Accordingly, we analyzed the dataset to determine if robust signal peptides could be identified that exhibited good performance across all single chain molecules. As shown in Figure 2, CHO homologous signal peptide E17 drove relatively high rates of ER translocation with all protein partners, facilitating 0.95-fold, 2.26-fold, and 2.61-fold increases in ETE IgG1 mAb LC, DTE IgG1 mAb LC and the ScFv fusion protein respectively, as compared to ISC. This novel element, derived from the CHO DDOST protein could replace incumbent signal peptides (such as ISC) in standard gene expression plasmids. While testing with a higher number of protein partners is required to definitively show generically robust performance across a wide range of product types, our data suggest that it could be deployed in single-chain protein-production vectors to deliver either (i) significant titer increases or (ii) similar effects to ISC.

### 2.3. Using machine learning to identify high-performance, molecule-specific signal peptide solutions *in silico*

While a toolbox approach permits identification of optimal signal peptides for a given protein, testing a large number of component combinations is not desirable in time (e.g., cell line development for biopharmaceutical protein production) and/or resource-limited contexts. Accordingly, we sought to develop a tool that could be utilized to screen signal peptide performance *in silico*, to minimize the required *in vitro* testing space while maximizing protein expression. While model-based tools have been created that can predict signal peptide performance in protein-partner specific contexts, this has only been achieved in bacterial systems (17). Utilizing the data obtained from screening our signal peptide panel in combination with three different molecules (i.e., Fig. 1), we attempted to build a model linking signal peptide performance to discrete protein sequence features. An XGBoosting (XGB) regression model was trained (Fig. 3) to predict recombinant protein titers as a function of eleven discrete sequence features, where sequences were defined as the relevant signal peptide in combination with the first 50 amino acids of the partner protein (33). The Input variables utilized were isoelectric point (pI), dipeptide stability, flexibility, aliphatic index, Gibbs free energy (∆G), grand average of hydropathicity index (GRAVY), and the percentage of glycine and proline residues in the signal peptide (GP%)). This feature set was designed to cover both physical (e.g., stoichiometry) and physiochemical (e.g., hydrophobicity) protein properties, whilst also incorporating specific characteristics that have previously been shown to effect signal peptide function (e.g., glycine/proline presence (34)).

**Figure 3:**
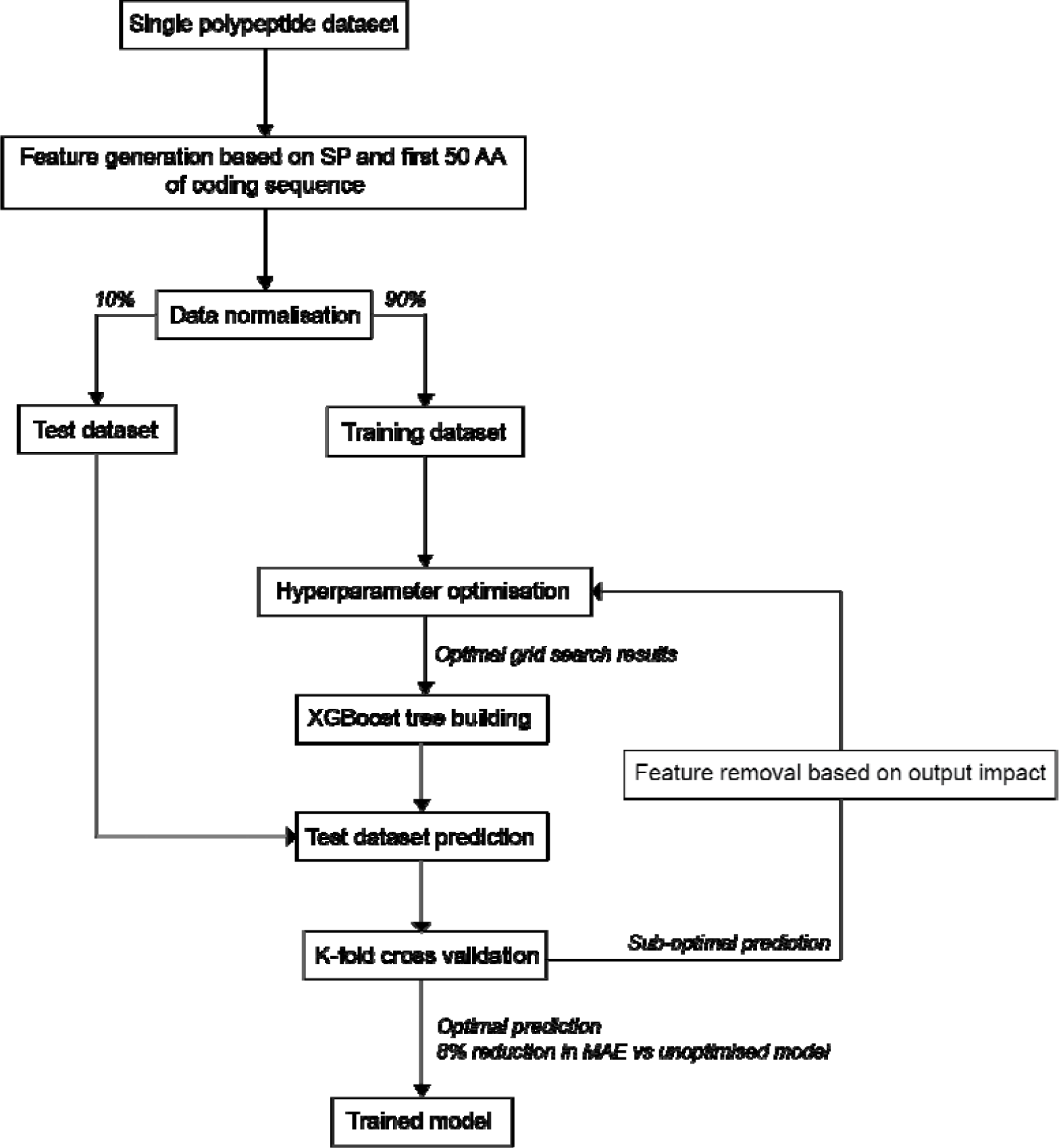
Schematic describing the creation of an XGBoost regression model for signal peptide selection in a molecular context. Using single chain molecule data described in Fig. 1 an XGBoost regression model was trained to predict signal peptide rank in combination with the first 50 amino acids of its partner protein. Features based on 7 protein parameters (isoelectric point (pI), dipeptide stability, flexibility, aliphatic index, Gibbs free energy (∆G), grand average of hydropathicity index (GRAVY) and the percentage of glycine and proline residues in the signal peptide (GP%)) were assigned to each signal peptide and its matching protein (a total of 114 combinations). All values were normalized using a min-max scale. Data was separated using a randomized 90%-10% train-test split. The model was optimized using a hyperparameter optimization grid search and employs early stopping to avoid overfitting. Optimized model K-fold cross validation mean absolute error (MAE) is 0.149 (0.023 SD).

Hyperparameter optimization was done using a grid search approach, and early stopping was employed to avoid model overfitting (model parameters and feature generation is described in detail in Section 4.5). An optimized model, where 7/14 features had a significant impact on predicting signal peptide performance, was moderately accurate in predicting the activity of a withheld test dataset (R^2^ = 0.65, Fig. 4A). This represents the first regression model that can accurately explain the function of mammalian signal peptides across varying protein partners. K-fold cross validation mean absolute error of the optimized model was 0.149 (0.02SD), an 8% decrease compared to the unoptimized model, confirming model robustness. We note that the predictive power of the model decreases as signal peptide activity (i.e., encoded ER translocation rate) increases. The Shapley additive explanation value for each training datapoint shows the relative impact of each sequence feature on model output (i.e., titer; Fig. 4B). Signal peptide activity was determined to be a function of both physiochemical (GRAVY, pI, dipeptide stability, ∆G) and stoichiometric (aliphatic index, flexibility) properties, where sequence pI had the greatest influence on construct activity. Although sequence pI correlates well with signal peptide activity in our model, it is unlikely to be a generically good predictor of element performance, which instead is determined by a complex interplay of multiple sequence features.

**Figure 4:**
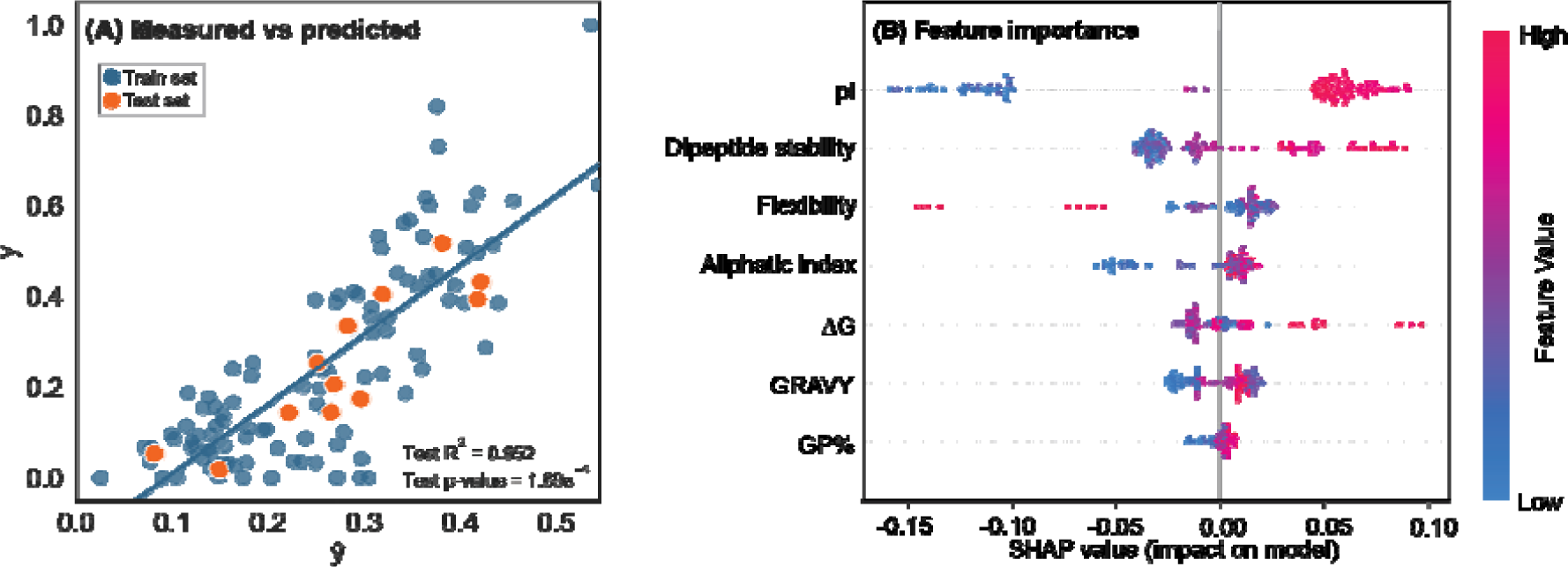
Graphical representation of model fit and the importance of relative features. (A) Moderate correlation is seen between measured and predicted ranking of the withheld test dataset (orange marker). Train dataset R^2^ = 0.772, p-value = 1.98e^-27^, confidence interval set at 95% (blue markers, blue line). (B) Individual feature importance SHAP values show input effects on model output. Positive SHAP values show a positive outcome, leading the model to predict a higher signal peptide ranking in its relative molecular context (high pI values results in higher ranking). Negative SHAP values show a negative outcome, leading the model to predict a lower signal peptide ranking in its relative molecular context (low pI values result in lower ranking). Each point represents one signal peptide in its molecular context. Features listed are isoelectric point (pI), dipeptide stability, flexibility, aliphatic index, Gibbs free energy (∆G), grand average of hydropathicity index (GRAVY) and the percentage of glycine and proline residues in the signal peptide (GP%).

Our intended use of this model was to (i) rationally select purely synthetic signal peptides able to increase expression of a given recombinant protein and (ii) significantly reduce the number of signal peptide constructs required to be made and tested. With respect to the latter for example, the data in Figures 1A-C (which largely results from the use of ‘natural” signal peptides) are derived from extensive informatic analyses to select context-relevant signal peptides – and the probability that another natural, signal peptide informatically-derived *de novo* would significantly increase the expression of a particular protein (e.g., over the ISC control) is approximately 30%.

Accordingly, a protein-specific synthetic signal peptide selection workflow was developed which utilized the same synthetic signal peptide library described in Section 4.1. which incorporated the use of SignalP6.0 to select high probability signal peptide sequences for subsequent model processing (Fig. 5A). We used the ScFv fusion protein experimental system (Fig. 1C) to test the utility of this approach against the criteria indicated above.

**Figure 5:**
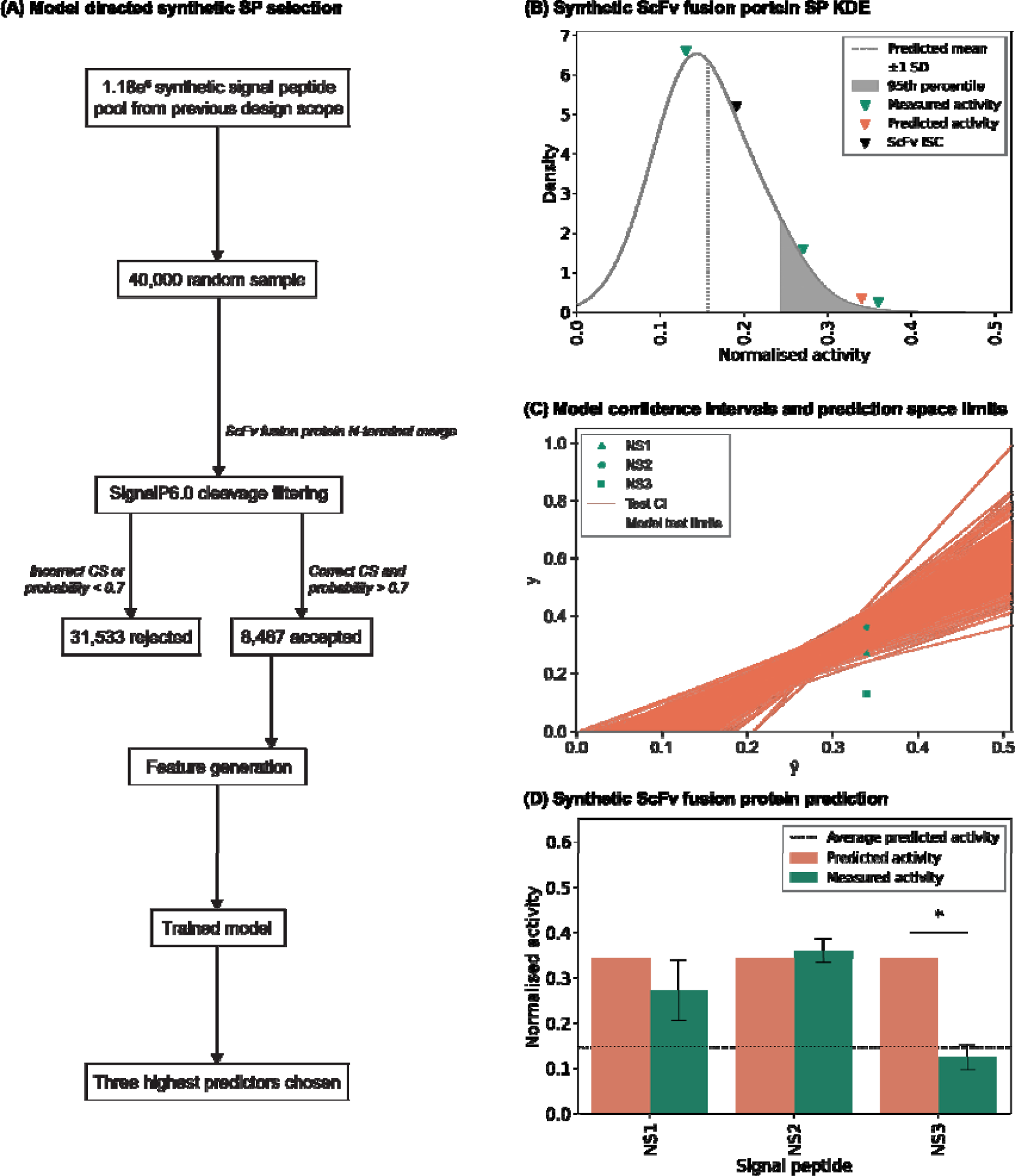
Functional performance of model derived synthetic signal peptides. Utilizing the model described in Fig. 3, a minimal test set of three predicted high activity synthetic signal peptides were selected for ScFv fusion protein expression from a random sample of 40,000 elements (A). The three synthetic signal peptides were selected from the 95^th^ percentile of 8,467 predicted synthetic signal peptides (B). Two of the three model chosen synthetic signal peptides showed measured activity which fell into the 95% confidence interval of each prediction, denoted by orange lines (C). Test signal peptides described in Fig. 4A defined the model prediction space, shaded in grey. The predicted activity of each synthetic signal peptide was directly compared to their measured activity counterpart, highlighting the acceptable predictability of two of the three synthetic elements in their ScFv fusion protein context (D). The ScFv fusion protein was independently transfected into CHO-K1 derived cells followed by measurement of secreted recombinant protein titer after 5d culture. NS denotes new synthetic signal peptides; refer to Table 2. Experimental data was normalized with respect to the mean volumetric titer observed on transfection of the ScFv fusion protein ISC then were then appropriately scaled for comparison. The mean value of all predicted synthetic signal peptides is represented by the dotted line. Each measured activity bar (green) shows the mean ± SD derived from three independent transfections, each performed in duplicate. An asterisk represents signal peptides which fall outside of the 95% confidence interval of model prediction.

A random sample of 40,000 synthetic elements was reduced to 8,467 which adhered to SignalP6.0 defined correct cleavage and had a signal peptide probability of >0.7. Following sequence feature generation, the performance of each signal peptide was predicted using the developed model. Based on the model’s moderate predictive power (R^2^ = 0.65), we rationalized that we could reduce the number of synthetic signal peptides to be tested to a minimal panel of three discrete peptides (Table 2) which were all hypothesized to yield ScFv fusion protein expression in the 95^th^ percentile of all predicted synthetic signal peptides (Fig. 5B). Model selected synthetic signal peptide-ScFv fusion constructs were evaluated in a 5-day fed batch transient production process. Of the three signal peptides chosen, two (NS1, NS2) facilitated significantly enhanced titers in comparison to ISC, where the best performing construct increased product yield by 1.95-fold. NS1 and NS2 also exhibited correlation between their predicted and measured activities, with acceptable prediction being defined as falling within the 95% confidence interval of experimental activity (Fig. 5C-D). In accordance with the model R^2^, two of the chosen predicted high activity synthetic elements performed as expected when expressing the ScFv fusion protein. Our data therefore confirm that model-based synthetic signal peptide selection is feasible, to reduce an impractical testing sample size substantially. Put in context, and in comparison with this model-based approach, the probability that any two additional informatically-derived signal peptides would both significantly increase expression of the test ScFv fusion protein (Fig. 1C) is approximately 10%.

**Table 2:**
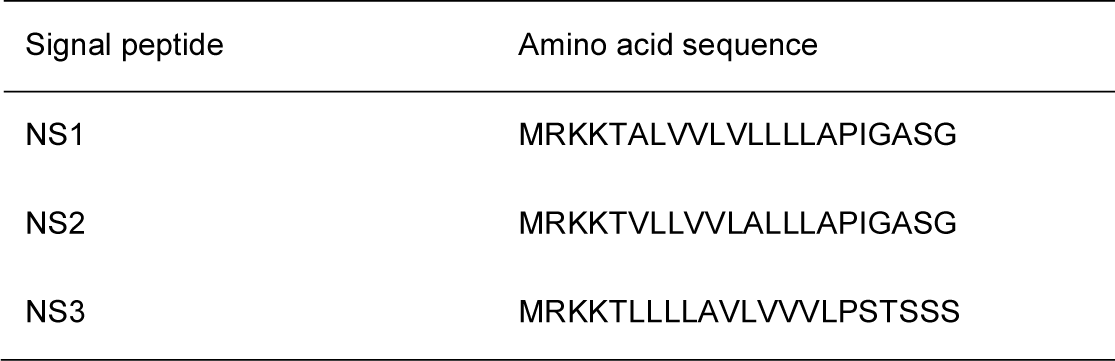
Model directed selection of synthetic signal peptides for in vitro testing with an ScFv fusion protein. Signal peptide ‘C’ is used as an ISC; refer to Table 1. Synthetic signal peptides were taken from initial design space pool as described in Section 4.1

| Signal peptide | Amino acid sequence |
| --- | --- |
| NS1 | MRKKTALVVLVLLLLAPIGASG |
| NS2 | MRKKTVLLVVLALLLAPIGASG |
| NS3 | MRKKTLLLLAVLVVLPSTSSS |

Mammalian signal peptides were also selected based on model prediction; however, prediction error was greater when compared with the synthetic signal peptides chosen (Fig. S1, Table S1). As discussed previously, the model’s predictive power decreases with increasing signal peptide activity, which may account for 2 out of three mammalian signal peptides exhibiting unpredictable functionalities *in vitro* (NE1, NE2). Though the model prediction threshold is theoretically limitless it is unreasonable to assume that this could translate to a biological system. In contrast to the predicted synthetic signal peptide choices, the predicted mammalian signal peptides fell outside of the known data limits of the model, further contributing to unpredictable functionalities in ScFv fusion protein expression (Fig. S1B).

We did not apply sequence homology-restrictions when selecting the *in vitro* testing panel. Accordingly, the synthetic designed elements that were selected shared similarities in their amino acid compositions, for example all having the same N-domain sequence. Indeed, NS1 and NS2 had identical amino acid compositions arranged in different discrete orders. Five H-domain amino acid rearrangements (position 6: A ➔ V, position 8: V ➔ L, position 10-12: LVL ➔ VLA) are sufficient to increase NS2 activity in comparison to NS1. This highlights a limitation of the model, which does not consider relative amino acid order when generating sequence features. We hypothesize that future models, utilizing larger datasets, that are able to include amino acid order (particularly in the H-domain) as an input parameter will have enhanced predictive power. However, despite only considering broader, overall sequence properties, we were able to build a model that substantially reduced the *in vitro* testing space required to identify a high-performing signal peptide for a specific protein-partner. For single chain proteins this can be utilized to either (i) significantly increase vector optimization studies by selecting a minimal subset from the original 37 component library and/or (ii) significantly increase the design space, permitting identification of context-specific high activity elements from large signal peptide databases (e.g., large synthetic libraries). To fully validate the utility of this approach, future studies will need to apply the model to a large panel of new protein-partner molecules.

### 2.4 Signal peptide engineering significantly enhances mAb production titers, where optimal vector designs are highly molecule-specific

Having validated the performance of our signal peptide library to enhance expression of single chain molecules, we next evaluated its utility to optimize production of more complex, multi-chain proteins. Monoclonal antibodies are the dominant class of biopharmaceutical product (1, 35), where both the ‘ideal’ HC:LC expression ratio, and the optimal absolute expression level of each chain, are highly molecule-specific (7, 9). Accordingly, a universal signal peptide combination will not facilitate maximal titers across product portfolios, necessitating screening to identify mAb-specific solutions. However, even for 2-chain molecules, a full-factorial analysis of all signal peptide combinations in our toolbox would entail 1369 permutations. Given that this screening burden would be intractable in most contexts, we concluded that a two-step optimization process would facilitate efficient derivation of optimal signal peptide combinations, where all 37 elements are first tested in association with the LC to identify parts that facilitate low, medium and high rates of expression (LC preferred over HC as it secreted when expressed in isolation, permitting rapid titer quantification). Restricting the number of options for the LC expression parameter to three experimentally-verified levels (while maintaining all 37 potential expression values for HC expression) a fractional factorial analysis of all part combinations requires 111 unique permutations, reducing the testing space by > 90%.

Using data from our screen of signal peptides in association with two discrete LC molecules (Fig. 1), we designated appropriate parts as driving high (maximum fold change in titer relative to ISC), medium (equivalent expression to ISC) or low (0.8-fold titer compared to ISC) activity components. A single element facilitating each discrete expression level was selected for the ETE (X7 > E17 > X4) and DTE (E1 > X8 > E3) LC. These components were utilized to drive ER translocation of the LC, where HC translocation rate was controlled by one of the 37 elements from the larger signal peptide library (i.e., 111 part combinations for each mAb). Given that it is common industrial practice to utilize different signal peptides for each protein chain, for each mAb we used a reference dual control system (RDCS) comprising ISC (driving HC translocation) in combination with an element of equivalent experimentally-verified strength (E17-LC and X8-LC for the ETE and DTE products respectively; Fig. 1). HC and LC expression vectors were transiently co-transfected into CHO cells and relative product titers were determined by ValitaTitre after a 5-day fed batch production process (Fig. 6A, 6C). Signal peptide assemblies resulted in diverse IgG1 titer outputs, ranging from 0.36-fold (LC:X7, HC:E5) to 1.82-fold (X7:X3), and 0.1-fold (E1:E6) to 2.12-fold (E3:E10) for the ETE and DTE mAb respectively, as compared to RDCS. The utility of the signal peptide toolkit-approach was validated by identification of least 26 element combinations that outperformed RDCS for each mAb. Indeed, ∼25% of part-assemblies tested facilitated significant increases in titer relative to RDSC, suggesting that the testing space required to identify vector engineering solutions could be substantially reduced.

**Figure 6:**
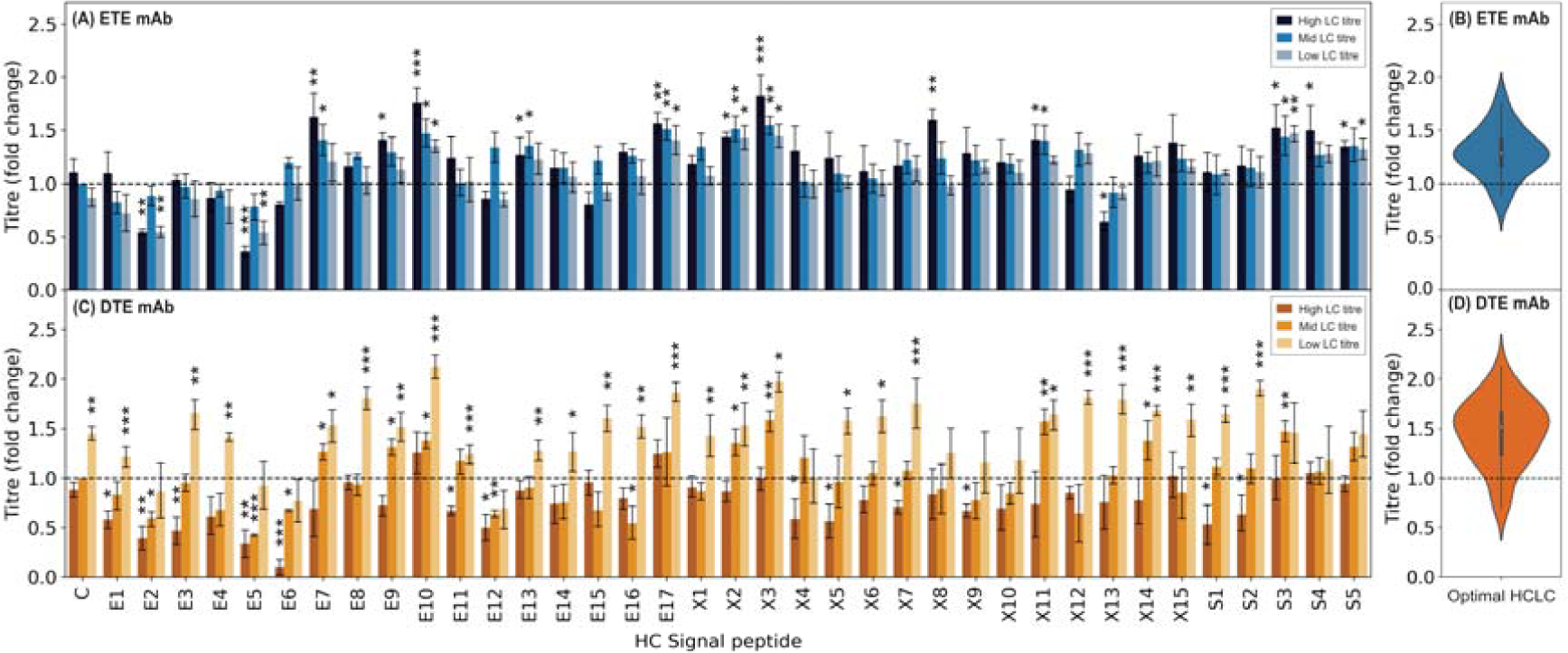
Choice of HC and LC signal peptide combinations significantly impacts both recombinant ETE IgG1 mAb and recombinant DTE IgG1 mAb production. Expression constructs (a total of 340 unique co-expression combinations) each encoding one of 37 mammalian signal peptides (Table 1) driving HC translocation with one of three recombinant mAb LC signal peptides (A, C) were independently transfected into CHO-K1 cells followed by measurement of secreted recombinant protein titer after 5d culture. (A) ETE IgG1 mAb, LC signal peptides are X7 (high), E17 (mid) and X4 (low); refer to Fig. 1A. (B) Maximized ETE IgG1 mAb recombinant protein volumetric titer distribution when one of 37 signal peptides expressing HC is combined with the optimal of three LC signal peptides expressing LC (X7, E17, X4; refer to Table 1). (C) DTE IgG1 mAb, LC signal peptides are E1 (high), X8 (mid) and E3 (low); refer to Figure 1B. (D) Maximized DTE IgG1 mAb recombinant protein volumetric titer distribution when one of 37 signal peptides (Table 1) expressing HC is combined with the optimal of three LC signal peptides expressing LC (E1, X8, E3; refer to Table 1). Data were normalized with respect to the mean volumetric titer observed on transfection of each mAb RDCS (ETE IgG1 mAb LC: MKMGVRLAARAWPLCGLLLAALGGVCA, DTE IgG1 mAb LC: MGSAALLLWVLLLWVPSSRA, HC: MGWSCIILFLVATATGVHS) (dotted line). Signal peptides are divided into three groups, E (CHO homologous, ETE: dark blue bars, DTE: brown bars), X (literature-mined, ETE: blue bars, DTE: orange bars) and S (synthetic, ETE: grey-blue bars, DTE: yellow bars); Table 1. Each bar shows the mean ± standard deviation derived from three independent transfections, each performed in duplicate. Statistical significance is defined as p ≤ 0.05 (* = p≤ 0.05, ** = p≤ 0.01, *** = p≤ 0.001).

**Figure 7.**
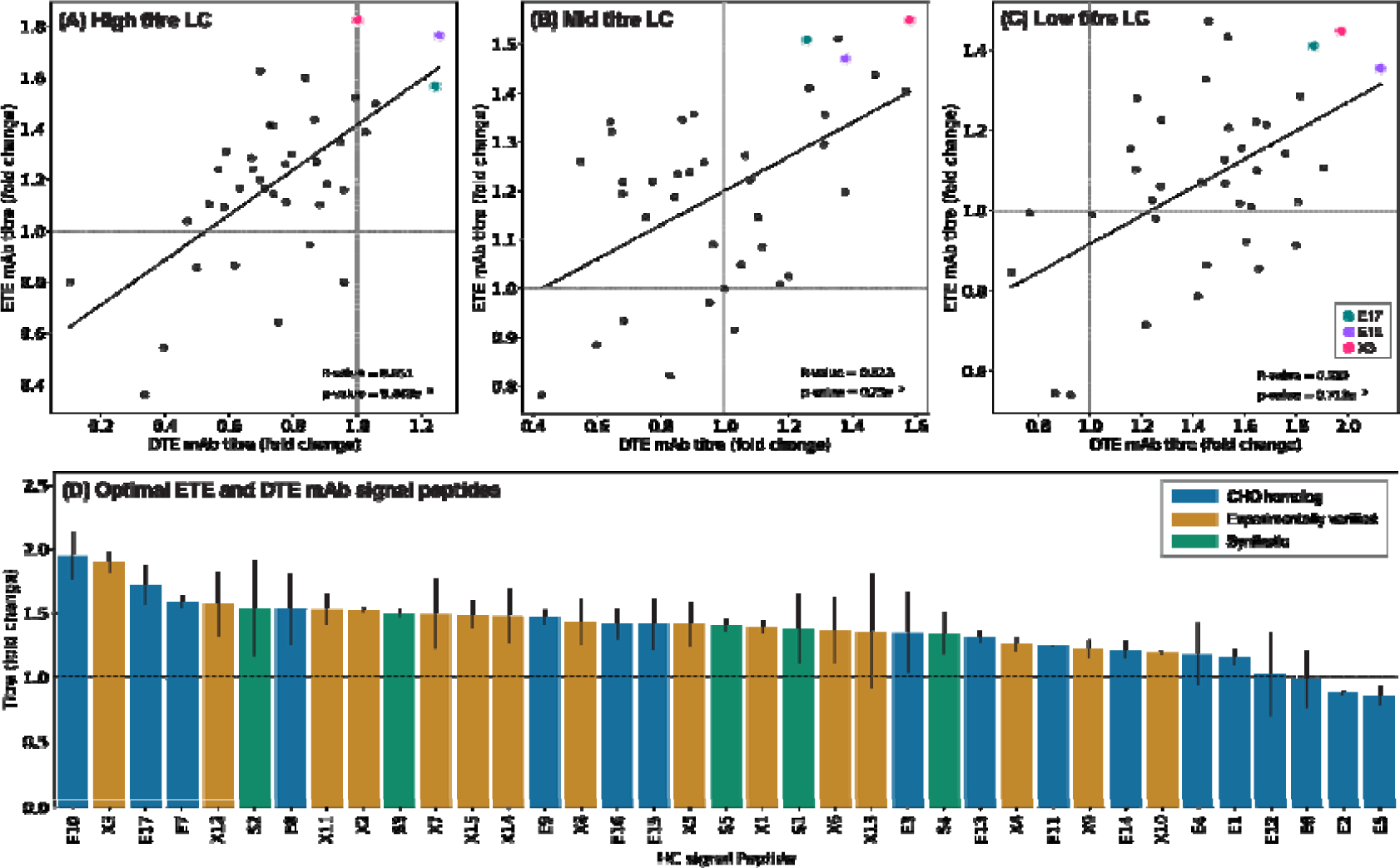
: Optimal LC signal peptide pairings in recombinant ETE IgG1 mAb and recombinant DTE IgG1 mAb identifies generic high performing HC signal peptides for recombinant protein production. Grouping ETE IgG1 mAb and DTE IgG1 mAb titer by respective LC titer (refer to Fig.6) shows moderate positive correlation between the ETE and DTE mAbs (A-C). Highlighted signal peptides show high volumetric titer across all combinations of LC (E10, E17, X3). Grey dashed line represents quadrant separation. Derived from the data shown in Fig. 6, a variety of CHO endogenous, literature-mined, and synthetic signal peptides (E17, E10, X3) yielded maximum volumetric titers (D). Bars represent the mean recombinant protein titer across two mAbs containing a mAb specific optimal LC signal peptide and one of 37 HC signal peptides (Fig.6). Data are normalized with respect to the RDCS (dotted line). Error bars represent the volumetric titer range across the optimal HCLC signal peptide combination for ETE IgG1 mAb and DTE IgG1 mAb tested for each HC signal peptide.

However, rules for designing smaller testing spaces are clearly molecule specific. For example, for the ETE protein, titers were generally enhanced by using the high strength signal peptide to control LC translocation. However, the inverse was true for the DTE mAb, where the low strength LC signal peptide outperformed the high and medium strength elements in the vast majority of vector designs tested, being the optimal partner for 35/37 HC-signal peptides. This may be explained by the proteins having contrasting optimal LC:HC expression ratios (7, 36), where enhancing DTE LC translocation rates may result in increased LC-aggregate formation (6). Alternatively, it could be a result of the proteins varying molecular structures. The ETE IgG1 mAb contains a kappa (κ) LC whereas the DTE IgG1 mAb contains a lambda (λ) LC. Assembly of HC-LC intermediaries is slower in λLC mAbs compared with κLC mAbs due to differing relative disulphide bond positioning (37, 38). Accordingly, using a high-strength signal peptide to maximize ER translocation of the LC may result in dyssynchronous mAb assembly processes.

As shown in Fig. 6B and 6D, there was a clear difference in the average performance of optimal LC:HC signal peptide combinations between the ETE and DTE mAb. The median molecular titer of best-performing element assemblies (i.e., each HC signal peptide in combination with the LC signal peptide partner that facilitated highest product expression) was significantly higher for the DTE mAb (1.52-fold compared to RDCS), compared to the ETE (1.29-fold), with a concomitant increase in the interquartile range. We therefore concluded that tailoring polypeptide ER translocation rates had a larger relative impact on DTE product expression. This is likely to be the case across product portfolios, as molecules that have been designated DTE typically have biosynthetic pathways rates that are dyssynchronous with cellular capacities, leading to induction of internal stress response pathways. Accordingly, for DTE products, it is likely that minimized signal peptide testing spaces can be used to identify product-specific vector solutions that significantly increase protein titers.

The relative impact of utilizing a discrete signal peptide to control HC ER translocation rate was moderately consistent across both (i) variable LC signal peptide partners encoding low-high ER translocation rates and (ii) the two different mAb molecules (Fig. 7A-C). This is in contrast to the data observed when LCs were expressed in isolation, where relative signal peptide performance showed no correlation between different single chain proteins (Fig. 2A-C). This indicates that the utility of signal peptide components for controlling HC expression rates is less context-specific, where discrete parts may display reasonably consistent performance across varying product and vector designs. Indeed, as shown in Figure 7D, when LC translocation rate is optimized, the performance of 37 signal peptides driving HC expression is highly consistent across the two different mAbs. This raises the possibility of utilizing universal signal peptides, for example E10 and X3 were the top two ranked HC signal peptide components for both products (elements ranked by ability to increase product titer relative to RDCS). Future studies will test the performance of these elements across a wider panel of mAb products to evaluate if they can be used to generically enhance mAb titers irrespective of HC partner-context.

Ideally, the testing space would be refined *in silico* using an appropriate model to predict the effect of signal peptide combinations on expression of multi-chain proteins. However, whilst this was achieved for single chain molecules (Section 2.3), we did not anticipate it would be possible to utilize an XGB model to forward engineer optimal element assemblies that maximize mAb titers. Indeed, our data highlights the unpredictable parameters governing expression of multi-chain products, where optimal stoichiometric HC:LC expression ratios are dependent on a complex interplay between product assembly pathways and host-cell biosynthetic capacities. To confirm this, we applied our previously described model (trained using data from our single-chain protein expression screens) to retrospectively predict the ability of signal peptide combinations to enhance mAb titers. Sequence features were generated for signal peptides in association with LC and HC partners, where overall values for discrete element assemblies (e.g. E1-LC:E10-HC) were calculated as the mean of the two part-polypeptide combinations (e.g. pI = (pI of E1-LC + pI of E10-HC)/2). As expected, the model had poor predictive ability, with only ∼38% (14/37) and ∼46% (17/37) of predictions were correct for the DTE and ETE product respectively, where correct prediction was defined as being in the 95% confidence interval range of experimental results. Accordingly, we concluded that whilst multi-chain product testing spaces can be reduced by selecting a small number of LC signal peptides *in silico*, it is intractable to accurately predict the performance of complete signal peptide compositions, necessitating the *in vitro* testing of 10s of potential vector solutions.

## 3. CONCLUSION

We have created a panel of signal peptides that can be utilized to enhance expression of recombinant proteins in CHO cells, validating the utility of three distinct component design/selection strategies that can be applied to other cellular contexts. As with previous studies in mammalian cell systems, we found that optimal signal peptide solutions were highly protein-specific. However, for all products tested we were able to derive vector designs that enhanced product titers by >1.8-fold, compared to standard industry technologies. Moreover, for single-chain products, we were able to build an XGB model that could guide selection of context-specific high-performing synthetic signal peptide elements. This model can be utilized to significantly reduce the screening space required to identify high performing, product-specific signal peptide solutions, representing the first time that such a model has been developed for a mammalian cell context. Although *in silico/in vitro* screening is required to identify the optimal signal peptide element for a new single chain molecule, we identified a small number of constructs that exhibited robust performance across different protein partners. For time and/or cost sensitive applications, these ‘universal’ signal peptides could be used as a generic expression vector component.

As expected, modelling techniques could not be applied to multi-chain mAb proteins, owing to unpredictable, molecule-specific optimal LC:HC expression ratios that are a function of internal cellular capacities and protein assembly dynamics. Indeed, we showed for the first time that specifically slowing down LC ER translocation rate can increase production of a DTE mAb. Despite this unpredictability, we were able to significantly reduce the vector testing space required to identify signal peptide combinations that increase product titer. Pre-selection of LC signal peptides that encoded low, medium and high levels of ER translocation focused the testing space towards solutions that enhanced protein production. Accordingly, in this work we have presented novel signal peptide parts, with associated streamlined *in silico* and *in vitro* testing processes, that can be used to rapidly re-design expression vectors to improve production of both simple and complex protein products.

## 4. METHODS

### 4.1 Synthetic signal peptide creation

A library of 1,168 experimentally verified human and mouse signal peptides with an amino acid length of 15-30 were extracted from signalpeptide.de and used as building blocks for synthetic signal peptide creation. Each signal peptide was separated out into its constituent N-, H- and C-domains. The first amino acid and final three amino acids were designated as minimal N- and C-domains respectively. The H-domain was identified using a sliding window approach, where the first and last 6AA region containing at least four hydrophobic amino acids (F, I, W, L, V, M, A, Y, C) marked the beginning and end of the domain. The amino acid sequences either side of the identified H-domain were assumed to be in the N-domain or C-domain.

Domain boundaries were assigned according to the following rules:

i. The N-domain must start with M and has a maximum length of ≤ 10 amino acids. It is of variable length.
ii. The H-domain is composed of amino acid blocks of six where four of the six amino acids must be hydrophobic (F, I, W, L, V, M, A, Y, C) and two must be non-hydrophobic. The maximum H-domain length is 12 amino acids, the minimum H-domain length is 6 amino acids.
iii. The C-domain is of variable length. It has a minimum length of 3 amino acids and a maximum length of ≤ 10 amino acids.

Where possible signal peptide composition rules described in literature were applied to each domain. Excluding basic domain separation, the definitions of each domain are limited with the C-domain being the most investigated. The synthetic N-domain was purely composed of conserved amino acids present in human and mouse signal peptide N-domains and always started with M. Amino acid conservation in the N-domain of the selected human and mouse signal peptides showed low amino acid preference. An arbitrary cut-off of ≥ 20% was applied resulting in synthetic N-domain amino acid selection being limited to K, T, A, G, S, P and R residues at amino acid positions -21 to -18 (where the last signal peptide residue position is -1). The synthetic H-domain was composed of conserved amino acids present in ≥ 60% human and mouse H-domain signal peptides (L, A, V) with the exception of the last H-domain amino acid (position -6) which was limited to P or G residues (*39, 40*). Previously published A-X-B, no P residues and -3, -1 literature defined C-domain rules were applied to synthetic C-domain creation (14, 29, 30). These applied rules resulted in different amino acid choices at each C-domain position. All five C-domain positions could be composed of A, G or S residues. At position -5 L, V, and I residues could also be present. At position -4 T or C residues could be additionally present and at positions -2 and -1 C and T residues could be respectively present.

Domain amino acid permutations were done resulting in 2,401 synthetic N-domains, 354,294 H-domains and 1,440 C-domains. N-, H- and C-domain permutations resulted in 1.2e^12^ synthetic signal peptides. To reduce this number 1% of the most different domains were chosen for synthetic signal peptide creation (24 N-domain options, 3,542 H-domain options, 14 C-domain options), giving a final synthetic signal peptide permutation number of 1.18e^6^. Using a SignalP4.1 signal peptide probability (D-score) limit of ≥ 0.7, five synthetic signal peptides were randomly selected for testing (41).

### 4.2 Molecular cloning for recombinant protein vector construction

Parental expression vectors containing the coding sequence (CDS) of an ETE IgG1 mAb, a DTE IgG1 mAb and an ScFv fusion protein (AstraZeneca, UK) were used for constructing signal peptide varied plasmids for recombinant protein assays. For each mAb, separate HC and LC plasmids were provided. Q5 site-directed mutagenesis kits (New England Biolabs, UK) were used to insert one of 37 signal peptides directly upstream of each CDS, replacing the control murine Ig HC signal peptide (MGWSCIILFLVATATGVHS; (27)). Transfection-grade plasmid DNA was purified using the QIAGEN plasmid plus Midiprep kit (QIAGEN, USA).

### 4.3 Cell culture and transient transfection

CHO-K1 derived host cells (AstraZeneca, UK) were maintained in CD CHO medium (Thermo Fisher Scientific, USA) supplemented with 6mM L-Glutamine. Cultures were maintained at 37ºC, 5% CO_2_ with 240RPM orbital shaking. Cells were routinely sub-cultured at a seeding density of 0.2e^6^ cells mL^-1^. Cell viability and concentration was measured using a VI-CELL viability analyzer (Beckman-Coulter, USA).

Cells were transiently transfected in a 96 well Amaxa Nucleofector System (Lonza, Switzerland) following the manufacturer’s protocols. Transfected cells were cultured in 24 shallow-well plates (Corning, UK) containing CD CHO medium supplemented with 6mM L-Glutamine for 5 days at 37ºC with 5% CO_2_ at 240RPM orbital shaking. Cultures were fed with a 1:1 Efficient Feed A (Thermo Fisher Scientific, USA) and Efficient Feed B (Thermo Fisher Scientific, USA) on day 3. Transient transfections of both mAbs were done using separate HC and LC plasmids at 1:1.

### 4.4 Recombinant protein quantification

Cell culture medium was clarified by centrifugation. ETE mAb LC and DTE mAb LC was quantified using Kappa and Lambda Human Immunoglobulin Free LC (FLC) ELISAs (BioVendor, UK) following the manufacturer’s protocol. Both IgG1 mAbs and ScFv fusion protein titer were quantified using ValitaTitre (ValitaCell, Ireland). ValitaTitre measurements were done in accordance with the manufacturer’s protocol. Commercially available purified kappa IgG1 mAb (Merck, Germany) and lambda IgG1 mAb (Merck, Germany) were used for quantification of ETE IgG1 mAb and DTE IgG1 mAb respectively. Purified ScFv fusion protein (AstraZeneca, UK) was used for quantification of the ScFv fusion protein. All assays were read using a SpectraMax iD5 microplate reader (Molecular Devices, USA).

### 4.5 Model creation

The XGboost package was used for construction and training of the model proposed (42). Titer from recombinant single chain proteins was normalized using a min-max scalar. Each signal peptide was paired with the first 50 amino acids of its respective protein and assigned 7 protein parameter generated features (isoelectric point, dipeptide stability, flexibility, aliphatic index, GRAVY, ∆G and signal peptide percentage of glycine and proline). Processed data was split into 90%-10% train/test. Hyperparameter optimization resulted in the following XGB regression parameters: colsample_bytree: 0.7; learning_rate: 0.05; max_depth: 3; min_child_weight: 1; n_estimators: 100; objective: reg:squarederror; subsample = 1. Early stopping was applied based on log loss validation (early stopping rounds = 5). K-fold cross validation parameters were as follows: number of splits = 5, number of repeats = 10. Mammalian experimental validation signal peptides were collected from UniProt using the following search term: annotation:(type:signal length:[10 TO 30]) taxonomy:”Eukaryota [2759]” AND reviewed:yes.

## Supporting Information

- Workflow of XGBoosting model application for mammalian signal peptide selection in the expression of an ScFv fusion protein (Fig. S1) and model selected mammalian signal peptide amino acid sequences (Table S1)

## Author Contributions

P.O performed all experiments, analysed all data, and wrote the manuscript. R.M provided technical guidance and supervision. A.J.B. and D.C.J. supervised the project design and execution. All authors discussed and revised the content of the manuscript.

## Notes

The authors declare no competing financial interest.

