## Supplementary information for "Protein-specific signal peptides for mammalian vector engineering"

^‡^SynGenSys Limited, Freeths LLP, Norfolk Street, Sheffield S1 2JE, U.K.

^§^Cell Line Development and Engineering, BioPharmaceuticals Development, R&D, AstraZeneca,

Cambridge, UK.

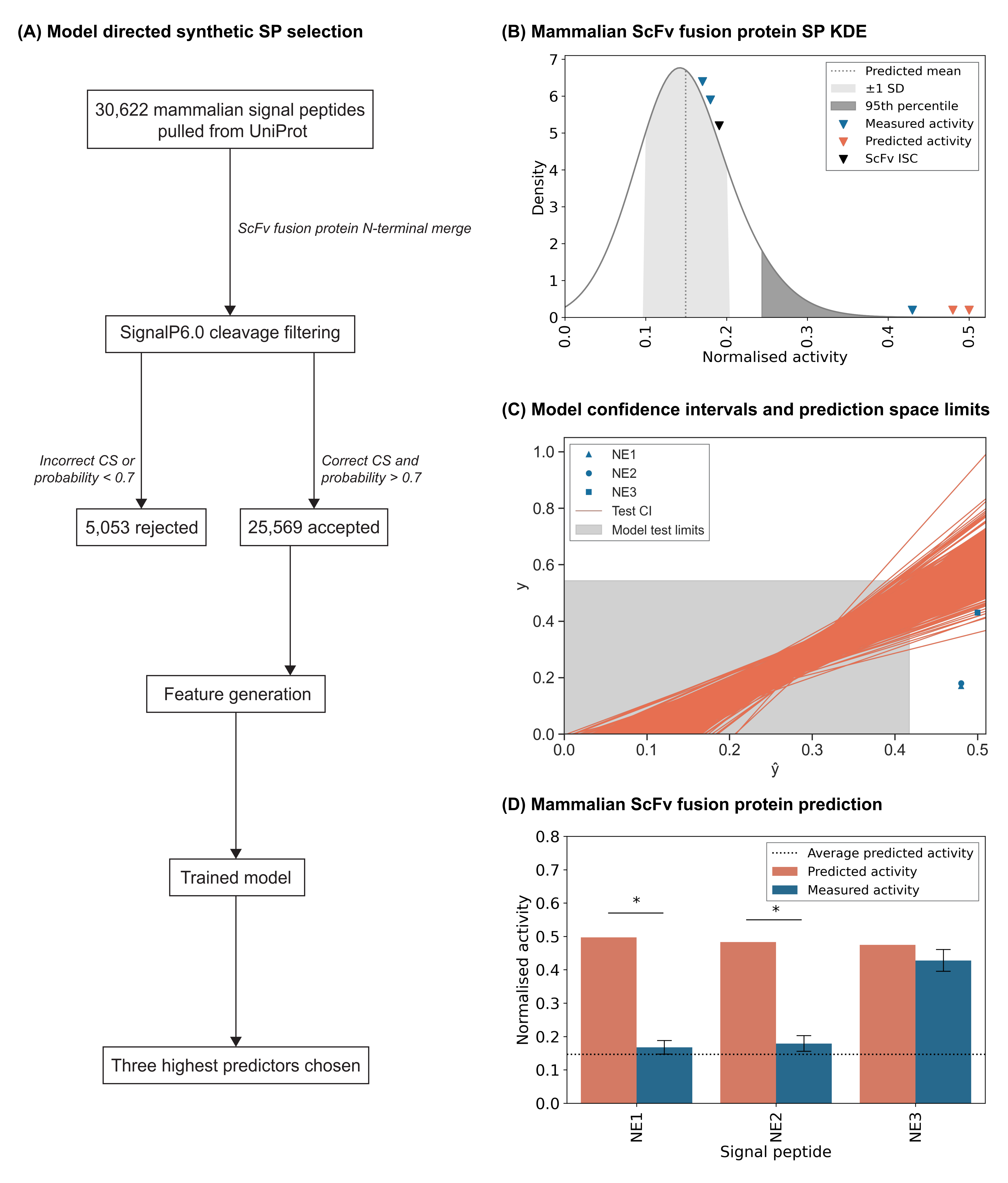

**Figure S1: Functional performance of model derived mammalian signal peptides.** Utilizing the model described in Fig.3, a minimal test set of three predicted high activity mammalian signal peptides were selected for ScFv fusion protein expression from 30,622 experimentally confirmed mammalian signal peptides; refer to Section 4.5) (A). The three mammalian signal peptides were selected from the 99^th^ percentile of 25,569 predicted signal peptides (B). One of the three model chosen mammalian signal peptides showed measured activity which fell into the 95% confidence interval of each prediction, denoted by orange lines (C). No chosen predicted mammalian signal peptides fell into the defined model prediction space, shaded in grey; refer to Fig.4A. The increase in error seen in mammalian signal peptides is likely a combination of increased prediction error with increased activity and mammalian prediction falling outside the validated model prediction range. The predicted activity of each mammalian signal peptide was directly compared to their measured activity counterpart, highlighting the acceptable predictability of one of the three endogenous mammalian elements in their ScFv fusion protein context (D). The ScFv fusion protein was independently transfected into CHO-K1 derived cells followed by measurement of secreted recombinant protein titer after 5d culture. NE denotes new mammalian signal peptides; refer to Table S1. Experimental data was normalized with respect to the mean volumetric titer observed on transfection of the ScFv fusion protein ISC then were then appropriately scaled for comparison. The mean value of all predicted mammalian signal peptides is represented by the dotted line. Each measured activity bar (blue) shows the mean ± SD derived from three independent transfections, each performed in duplicate. An asterisk represents signal peptides which fall outside of the 95% confidence interval of model prediction.

**Table S1: Model directed selection of mammalian signal peptides for in vitro testing with an ScFv fusion protein.** Signal peptide ‘C’ is used as an industrially relevant standard reference signal peptide (ISC); refer to Table 1.

| Signal peptide | Amino acid sequence | Signal peptide origin |
| --- | --- | --- |
| NE1 | MARGSLRRLLRLLVLGLWLALLRSVAG | N-terminal signal peptide of human tumour necrosis factor receptor superfamily member 12A. |
| NE2 | MARRSRHRLLLLLLRYLVVALGYHKAYG | N-terminal signal peptide of human junctional adhesion molecule B. |
| NE3 | MGTVRSRRLWWPLPLLLLLLRGPAGARA | N-terminal signal peptide of black-handed spider monkey proprotein convertase subtilisin/kexin type 9. |
